# Individualized Assays of Temporal Coding in the Ascending Human Auditory System

**DOI:** 10.1101/2021.09.13.460174

**Authors:** Agudemu Borjigin, Alexandra R. Hustedt-Mai, Hari M. Bharadwaj

## Abstract

Neural phase-locking to temporal fluctuations is a fundamental and unique mechanism by which acoustic information is encoded by the auditory system. The perceptual role of this metabolically expensive mechanism, the neural phase-locking to temporal fine structure (TFS) in particular, is debated. Although hypothesized, it is unclear if auditory perceptual deficits in certain clinical populations are attributable to deficits in TFS coding. Efforts to uncover the role of TFS have been impeded by the fact that there are no established assays for quantifying the fidelity of TFS coding at the individual level. While many candidates have been proposed, for an assay to be useful, it should not only intrinsically depend on TFS coding, but should also have the property that individual differences in the assay reflect TFS coding *per se* over and beyond other sources of variance. Here, we evaluate a range of behavioral and electroencephalogram (EEG)-based measures as candidate individualized measures of TFS sensitivity. Our comparisons of behavioral and EEG-based metrics suggest that extraneous variables dominate both behavioral scores and EEG amplitude metrics, rendering them ineffective. After adjusting behavioral scores using lapse rates, and extracting latency or percent-growth metrics from EEG, interaural timing sensitivity measures exhibit robust behavior-EEG correlations. Together with the fact that unambiguous theoretical links can be made relating binaural measures and phase-locking to TFS, our results suggest that these “adjusted” binaural assays may be well-suited for quantifying individual TFS processing.

## 1 Introduction

All acoustic information we receive is conveyed through the firing rate and/or timing of the neural spikes (i.e., rate-place vs temporal coding) of cochlear neurons. Temporal information in the basilar-membrane vibrations consists of cycle-by-cycle variations in phase—the temporal fine structure (TFS), and dynamic variations in amplitude—the envelope (ENV) (Hilbert, 1906). Cochlear neurons phase-lock to both TFS (Johnson, 1980), and ENV (Joris and Yin, 1992) robustly, with TFS phase-locking extending at least up to 1000 Hz (Verschooten et al., 2019). While the peripheral rate-place code has consistent counterparts throughout the auditory system, the upper limit of phase-locking progressively shifts to lower frequencies along the ascending pathway (Joris et al., 2004). How this metabolically expensive initial/peripheral temporal code (Laughlin et al., 1998; Hasenstaub et al., 2010) contributes to everyday hearing, and how its degradation contributes to perceptual deficits, are foundational questions in auditory neuroscience and clinical audiology. Yet, the significance of TFS coding is debated (Oxenham, 2013; Drullman, 1995; Swaminathan and Heinz, 2012; Oxenham and Simonson, 2009).

Previous studies have explored whether sound localization and pitch perception benefit from TFS cues. While it is established that lateralization of low-frequency sounds depends on TFS (Smith et al., 2002; Yin and Chan, 1990), whether TFS is important for pitch perception is difficult to ascertain. Behavioral studies suggest that low-frequency periodic sounds elicit a stronger a pitch than high-frequency sounds (Moore, 1973; Houtsma and Smurzynski, 1990; Bernstein and Oxenham, 2003), suggesting a possible role for TFS. However, these results permit alternate interpretations in terms of place coding and haromonic resolvability (Oxenham, 2012). Regardless of its role in quiet, whether TFS is important for masking release in noise is further debated, especially when other redundant cues can also convey pitch or location, and when room reverberation can degrade temporal cues (Best et al., 2005; Oxenham and Simonson, 2009; Ihlefeld and Shinn-Cunningham, 2011).

To investigate the role of TFS, studies have used sub-band vocoding to independently manipulate ENV and TFS cues (Smith et al., 2002; Hopkins et al., 2008; Hopkins and Moore, 2009; Lorenzi et al., 2009; Ardoint and Lorenzi, 2010). However, acoustic manipulations cannot eliminate subsequent confounding of ENV, TFS, and place cues without detailed knowledge of cochlear processing at the individual level (Oxenham, 2013; Swaminathan and Heinz, 2012). Thus, establishing the precise role of TFS through vocoding experiments is difficult, although the use of high-fidelity vocoders can help (Viswanathan et al., 2021). An alternative approach is to directly measure TFS sensitivity from individual listeners and compare it to individual differences in other perceptual measures. This approach has been successfully used to address other fundamental questions (McDermott et al., 2010; Bharadwaj et al., 2015; Whiteford et al., 2020). Unfortunately, the lack of established measures of TFS sensitivity at the individual level limits this enterprise.

Conventional behavioral TFS-sensitivity measurements have attempted to eliminate confounding cues such that primary task would rely on TFS processing (Strelcyk and Dau, 2009; Moore and Sek, 2009; Hopkins and Moore, 2010; Sek and Moore, 2012). However, they did not assess the influence of extraneous factors on the measured scores. Unfortunately, non-sensory factors can contribute significantly to individual variability even when the tasks themselves rely on specific acoustic cues (Kidd et al., 2007). Objective electrophysiological measures of TFS sensitivity can circumnavigate this problem; however, such studies are scarce (Verschooten et al., 2015; Parthasarathy et al., 2020). Here, we employ a battery of both behavioral and electroencephalography (EEG)-based measures of TFS sensitivity on a cohort of normal-hearing individuals to identify candidate assays of TFS processing at the individual level. Our results suggest that extraneous variables dominate both behavioral and raw EEG measures. However, with adjustments, we observed robust behavior-EEG correlations in binaural assays, rendering them well-suited for quantifying individual TFS processing.

## 2 Materials and Methods

The primary goal of the current study was to evaluate an array of both behavioral and electrophysiological measures as candidate assays of TFS sensitivity at the individual level. Based on the finding that nonsensory factors contribute significantly to behavioral TFS measures, a large-N supplementary behavioral experiment was conducted to assess whether non-sensory factors also influence ENV sensitivity when measured from naïve participants.

### 2.1 Participants

One hundred and fifty-three listeners, aged 18-60 years, were recruited from the local community near Purdue University. All human subject measures were conducted following protocols approved by the Purdue University Internal Review Board and the Human Research Protection Program. Participants were recruited via posted flyers and bulletin-board advertisements and provided informed consent. Of the 153 subjects, 44 (20 males) participated in the main experiments designed to evaluate candidate assays of TFS processing. The remaining N=109 participated in the supplementary experiment aimed at testing whether non-sensory factors also influence ENV sensitivity. Although the goal of the main experiment was to conduct all behavioral and electrophysiological TFS measures on each participant, some were not able to finish the full study battery due to limited availability.

### 2.2 Experimental Design and Statistical Analyses

#### 2.2.1 Behavioral Measures of the TFS Coding

Each of the following behavioral measurements was conducted on a different day from the others to randomize the influence of factors that may be idiosyncratic to a specific test day/session. A single lab visit contained only one behavioral measurement to reduce the impact of cognitive fatigue due to hour-long experiments.

##### Frequency Modulation (FM) Detection Thresholds

To obtain monaural TFS sensitivity, FM thresholds were measured separately in each ear, using a weighted (3:1) one-down-one-up (Kaernbach, 1991), two-alternatives-forced-choice (2AFC) adaptive procedure. The stimulus in the target interval was a 500-ms-long 500-Hz tone with frequency modulation at a 2-Hz rate and variable depth. The reference interval was a 500 Hz pure tone. The inter-stimulus gap was 900 ms. The stimulus was ramped on and off with a rise/fall time of 5 ms to exclude audible transitions. The stimulus level was 70 dB SPL. The subjects were instructed to press a button to indicate the interval containing the FM. Each measurement block was terminated after 11 reversals and the median of all the reversals from the adaptive procedure was extracted as the threshold. Four blocks of measurements were obtained in each ear from each subject. Except for an additional “demo” block to orient the participants before the formal testing, there was no further training. Sennheiser HDA 300 over-the-ear headphones were used for stimulus delivery. The slow FM rate of 2 Hz was chosen because it is thought that TFS cues are used to detect FM at rates below about 10 Hz (Moore and Sek, 1996; Strelcyk and Dau, 2009). However, recent evidence suggests that this may not be the case (Whiteford et al., 2020). Nonetheless, given the large body of literature using and interpreting slow-FM detection as a measure of TFS sensitivity, we chose to include this in the battery of candidate measures.

##### Interaural Time Difference (ITD) Detection Thresholds

To obtain a binaural measure of TFS sensitivity, we measured ITD detection thresholds using a three-down-one-up, two-alternatives-forced-choice adaptive procedure. The stimulus consisted of two consecutive 400 ms-long, 500 Hz tone bursts with an ITD. The leading ear for the ITD was switched from the first burst to the second. The stimulus was ramped on and off with a rise/fall time of 20 ms to exclude audible transitions and to reduce reliance on onset ITDs. The stimuli were presented at 70 dB SPL. Subjects were asked to report the direction of the jump (left-to-right or right-to-left) between the intervals through a button press. It was preferable to have subjects indicate the direction of change because absolute lateralization can be influenced by multiple factors (Moore and Sek, 2009). The threshold was defined as the geometric mean of the last nine reversals, and measured repeatedly across eight blocks, with a short break scheduled after the fourth block. Etymotic Research (ER-2) insert earphones were used for delivering the stimuli. A separate “demo” block was included before the experimental blocks to familiarize the subject with the task.

##### “Non-sensory” Score

Because the main goal of the study is to evaluate candidate measures of TFS coding in naïve subjects, i.e., individuals without extensive training/practice on the measured tasks, we anticipated that extraneous “non-sensory” variables may influence the measured thresholds. Accordingly, percent-correct scores on easy “catch” trials were calculated to quantify the subject’s engagement. Errors made in these catch trials reflect non-sensory factors such as lapses in attention, variations in motivation, alertness, etc. The criterion for designating a trial as a “catch” trial for the FM detection task was a frequency deviation (modulation depth) of 15 Hz, while it was 80 microseconds for the ITD task.

##### Supplementary amplitude modulation (AM) detection task

To further investigate the influence of non-sensory factors on behavioral measures in general, we carried out a supplementary experiment using a task that is unrelated to TFS processing—an AM detection task similar to (Bharadwaj et al., 2015). A similar 2AFC procedure as in the FM and ITD detection threshold measurements was employed. The target was a 500-Hz, 75 dB SPL band of noise centered at 4 or 8 kHz, and amplitude modulated at 19 Hz. Two unmodulated tones, flanked at two equivalent rectangular bandwidths (ERBs) (Glasberg and Moore, 1990; Moore, 1968) away from the center frequency, each at 75 dB SPL, were used to minimize off-frequency listening. The signal in the reference interval was statistically identical but unmodulated. Using a noise carrier helps eliminate spectral cues for the AM detection task (Viemeister, 1979). The threshold for the modulation depth detection was determined by an adaptive weighted one-up-one-down procedure (Kaernbach, 1991).

#### 2.2.2 Electrophysiological Measures of the TFS Coding

While behavioral measures directly assess perceptual sensitivity to TFS, they may also reflect common non-sensory factors such as attention and motivation. To dissociate TFS coding from non-sensory factors, we designed two passive EEG measures of the TFS coding and compared them to individual behavioral measures. Participants watched a silent, captioned video of their choice while passively listening to the auditory stimuli. EEG recordings were obtained using a 32-channel EEG system (Biosemi Active Two), while the stimuli were presented via ER-2 insert earphones.

##### Cortical correlates of TFS-based ITD processing

Cortical EEG was recorded in response to 70 dB-SPL 500-Hz tones that were amplitude-modulated (100% depth) at 40.8 Hz. 40.8 Hz can elicit a strong auditory steady-state response in EEG recordings (Picton et al., 2003); this response was used here as a measure of recording quality (Figure 1-B). As with the behavioral measurement, the leading ear for the ITD switched halfway through the trial. The ITD switch direction was randomized across trials. To minimize monaural cues, the ITD switch coincided with a trough of the 40.8 Hz modulation (Figure 1-A). The magnitude of the ITD was randomly chosen to be 20, 60, 180, or 540 *μs*. A total of 1200 trials were presented to the listener. The inter-stimulus interval was uniformly distributed between 500 and 600 ms. The recordings from the vertex electrodes, Cz and Fz, were used for analysis.

**Figure 1.**
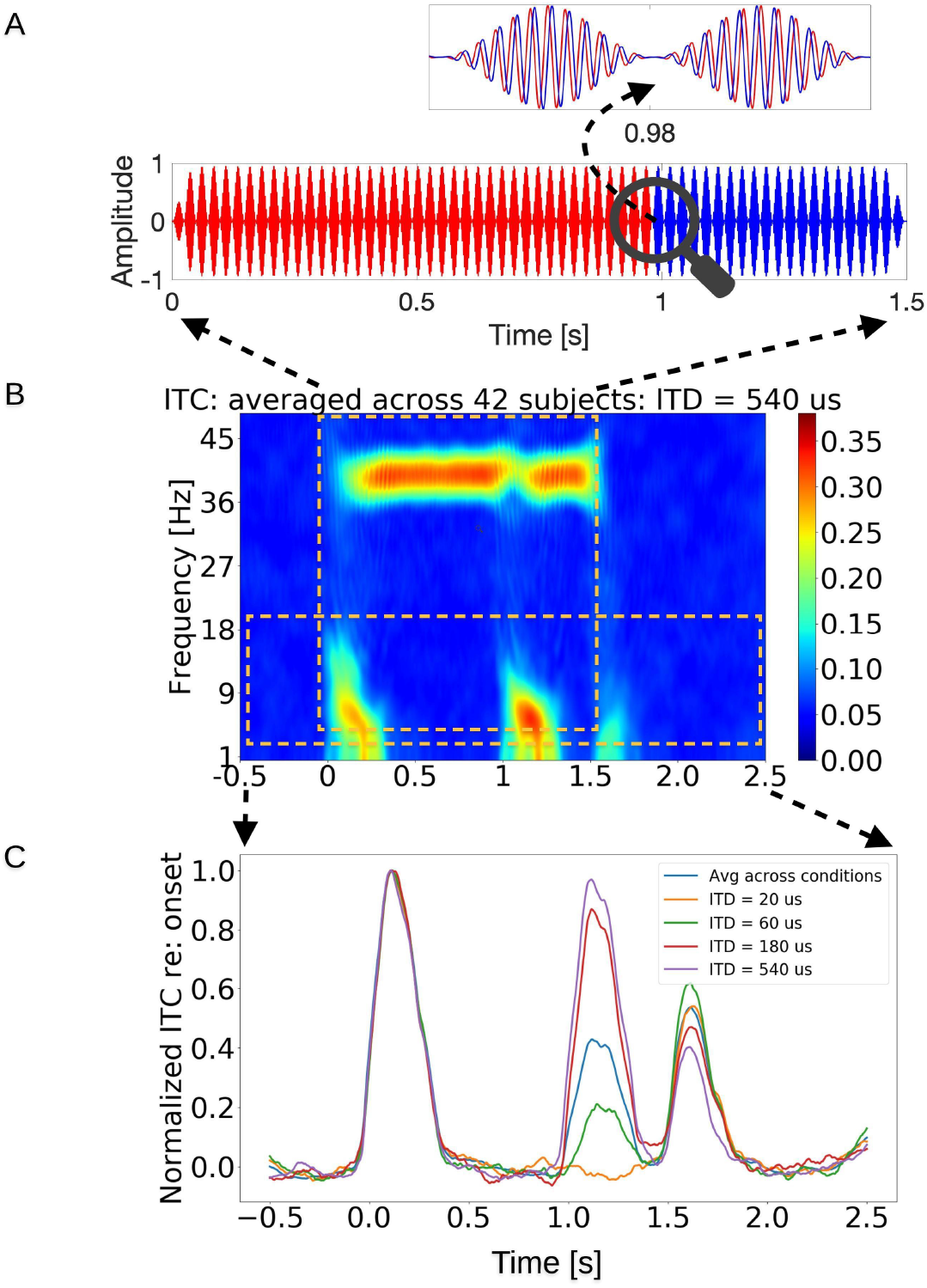
Stimulus paradigm and response from the EEG-TFS sensitivity measurement. A. The stimulus is a 1.5-second long, 500-Hz pure tone that is amplitude modulated at 40.8 Hz. The red color represents the sound leading in time in the right ear, whereas the blue stands for the sound leading in the left ear. The stimulus in the right ear leads in time till 0.98 seconds, after which the ITD shifts in polarity, i.e., the stimulus in the left ear takes the lead. The ITD jump occurs when the stimulus amplitude is zero to minimize the involvement of monaural cues. B. Spectrogram of the inter-trial coherence (ITC) of the EEG response, averaged across 42 subjects, with the colormap indicating the ITC. Robust auditory steady-state responses can be seen around the AM frequency of 40.8 Hz. There are also salient responses time locked to the stimulus onset, offset, and importantly, to the ITD jump. C. The average time course of the ITC for frequencies below 20 Hz is shown for each ITD jump condition. The response evoked by the shift in the ITD polarity increases monotonically with the size of the ITD jump confirming that the response is parametrically modulated by TFS-based processing.

##### Frequency Following Response (FFR)

Subcortical FFRs were measured in response to tones in a forward-masking stimulus configuration (Verschooten and Joris, 2014). The stimuli consisted of three consecutive segments: a 500-Hz probe tone that was 100-ms long and at 75 dB SPL, a “forward-masker” tone of the same frequency and duration but at 85 dB SPL, and the same probe tone. A 50-ms silent gap was included between the first probe tone and forward-masker, but only a 1-ms gap was included between the forward-masker and the second probe tone. Each stimulus segment was ramped on and off over 5 ms to reduce audible transitions. The polarity of the stimulus was alternated across a total of 8000 trials. The 500 Hz component of *differential* response obtained across the two stimulus polarities reflects response components that are phase-locked to the TFS, whereas the summed 500 Hz response represents the response to the ENV. However, the TFS component can contain both pre-neural (e.g., cochlear microphonic) as well as neural responses. Verschooten and Joris 2014 argued that the nonlinear residual obtained by subtracting the TFS response to the second probe tone from the TFS response to the first probe tone will isolate the neural component and suppress the approximately linear cochlear microphonic (CM). This is because the forward masking of response to the second probe tone only masks the neural component, whereas the CM is intact. Owing to the inner-hair-cell rectification, the summed response across the two polarities also contains a component at twice the stimulus frequency (1000 Hz) that reflects physiological currents phase-locked to the TFS in the stimulus. Although TFS-related, whether this double-frequency response is purely neural as has been previously interpreted (Parthasarathy et al., 2020), or whether it includes pre-neural contributions is unknown. Thus, we considered two candidate subcortical correlates of TFS processing: (1) The 500 Hz component derived from the differential response across the two polarities of stimulus presentation, and (2) the 1000 Hz component derived from the summed response across two polarities of stimulus presentation. Recordings from the vertex channels (i.e., Fz and Cz) were used for further data analysis.

#### 2.2.3 Statistical Analyses

Pearson correlations were calculated to illustrate simple associations between pairs of measurements. Statistical inference about behavior-physiology correlations was made using a multiple linear stepwise regression analysis by adding new potential predictors one by one to model the dependent variable. All reported significant associations met a false discovery rate criterion of 5% to control for multiple comparisons (Benjamini and Hochberg, 1995). Statistical analyses were performed using R (R Core Team, https://www.r-project.org/).

##### Code Accessibility

Stimulus generation and data analyses were done using custom scripts. They can be accessed at: https://github.com/AgudemuBorjigin/stimulus-TFS https://github.com/AgudemuBorjigin/EEGAnalysis, and https://github.com/AgudemuBorjigin/BehaviorDataAnalysis.

## 3 Results

### 3.1 Non-sensory factors contribute to large individual differences in behavioral measures of TFS coding

Similar to previous reports of large individual differences in the AM and ENV-based ITD detection thresholds across normal-hearing (NH) listeners (Bharadwaj et al., 2015), both the FM and TFS-based ITD detection thresholds varied widely across our NH listeners. FM detection thresholds across 43 NH listeners ranged from 7 to 22 dB relative to 500 Hz (i.e., a frequency deviation (Fdev) of 2-13 Hz from 500 Hz). ITD detection thresholds varied from 21 to 39 dB relative to 1 us (i.e., 11-89 us) across 37 NH listeners. These FM and ITD detection thresholds are shown along with the results from similar studies, in Figure 8 and Figure 9, respectively, and were largely comparable.

Across listeners, neither FM nor ITD thresholds (each averaged across repetitions and two ears) correlated with the audiograms; however, the two measures were significantly correlated with each other in a simple linear regression analysis (r = 0.44, p = 0.01, n = 33). While the correlations may arise from individual differences in TFS coding, they can also reflect non-sensory factors such as attention, motivation, etc. To disambiguate these competing explanations, we assigned each listener a “non-sensory score”, which was their lapse rate in catch trials (see Materials and Methods). When those scores were factored out from each measurement, the correlation between the monaural FM and binaural ITD thresholds dropped such that the association no longer met conventional statistical significance criteria (R = 0.31, P = 0.08, n = 33), suggesting that non-sensory factors play a large role in raw scores. Furthermore, when just the blocks with the largest (i.e., worst) FM and ITD thresholds for each subject were compared, considerably stronger correlations were observed (r = 0.6, p = 9e-4, n = 33), underscoring the involvement of non-sensory factors in behavioral measurements. Figure 2 shows the correlations between the measured and predicted thresholds solely based on the lapse rates (i.e., the non-sensory score). The involvement of non-sensory factors is evident, especially for the poorer performers.

**Figure 2.**
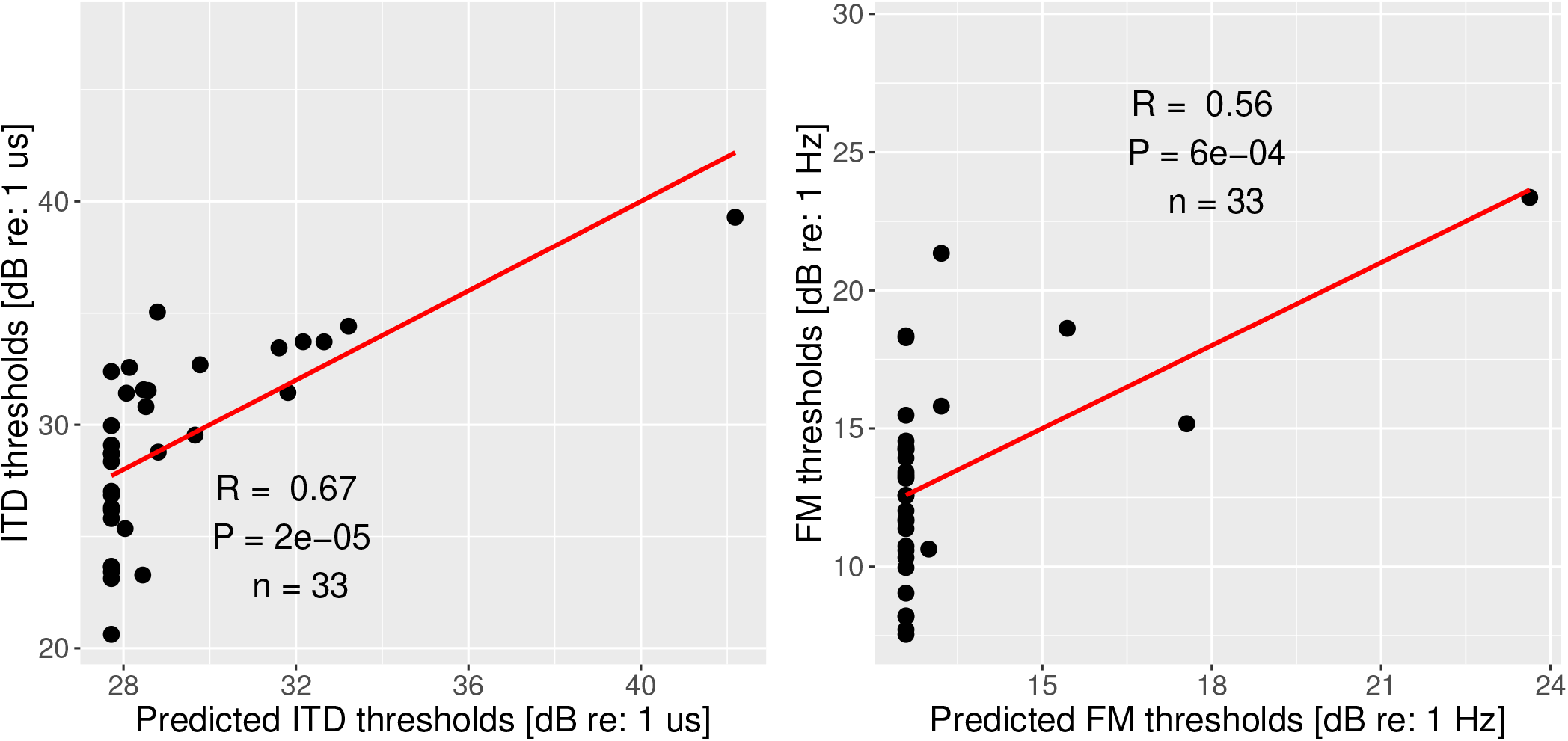
Measured vs. predicted thresholds based on lapse rate. [left] Measured vs predicted ITD detection thresholds; [right] Measured vs predicted FM detection thresholds. The significant contribution of non-sensory factors is apparent, especially for the poorer performers.

To confirm the involvement of non-sensory factors in raw behavioral scores, a similar comparison of thresholds and lapse rates was conducted for the supplementary AM detection task. The predicted thresholds based on the “non-sensory score” significantly correlated with the measured AM thresholds (R = 0.51, P = 9e-9, n = 108; Figure 3). This result indicates the significant weight of non-sensory factors, not only for FM and ITD detection measurements but behavioral measures in general.

**Figure 3.**
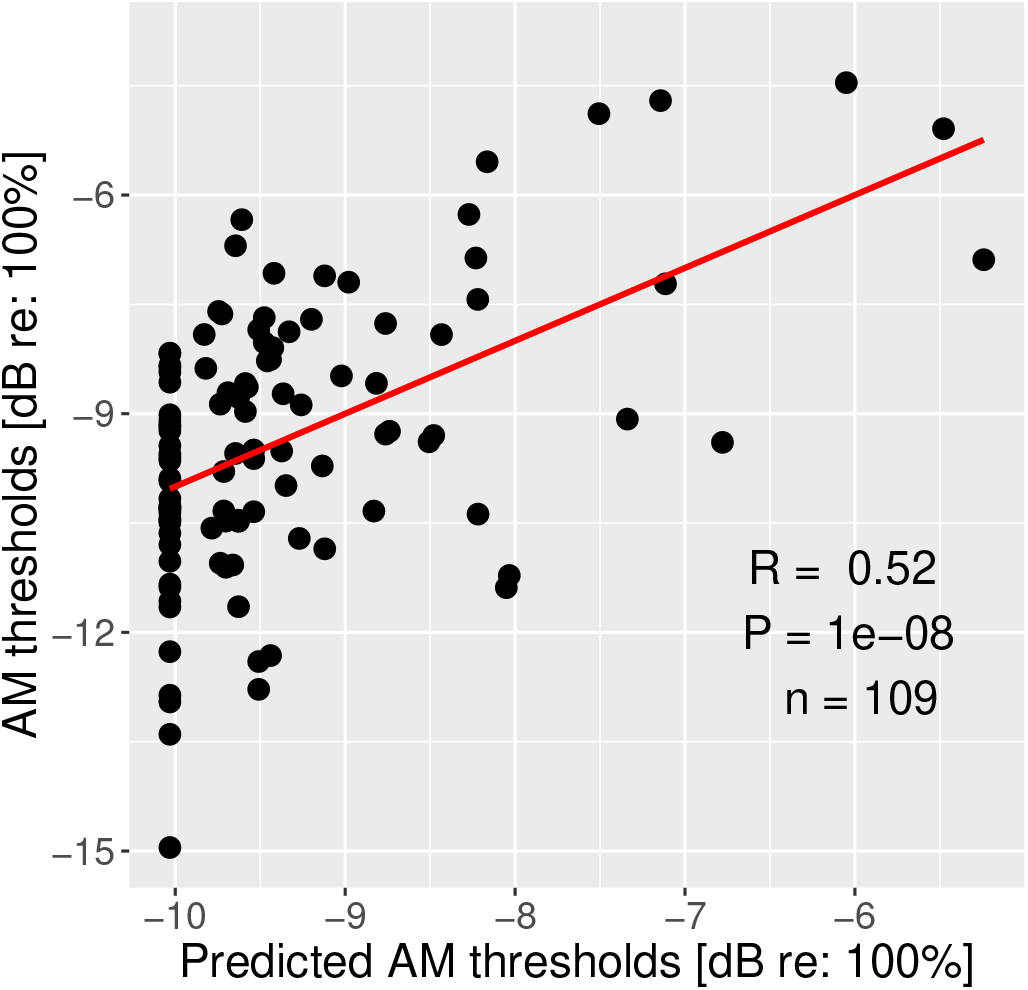
Measured vs predicted AM thresholds based on lapse rate. The thresholds are the average detection thresholds of AM tones at 4 kHz and 8 kHz. The significant significant contribution of non-sensory factors is apparent.

### 3.2 Raw electrophysiological TFS measures are strongly influenced by extraneous sources of variance

Two passive electrophysiological measurements were carried out to objectively evaluate individual TFS coding. Because passive electrophysiological measures are likely to be influenced by distinct extraneous factors (e.g., head size) compared to behavioral measures (e.g., motivation/engagement), these measurements provide a complementary window into individual TFS coding.

#### 3.2.1 Candidate cortical correlates of TFS processing

Cortical responses evoked by the polarity shift of the ITD are quantified through the phase-locking strength shown in the phase-locking spectrograms (Figure 1 B). Clear responses to the onset, offset, and ITD jump are apparent in the low-frequency portion of the phase-locking spectrogram. The sustained auditory steady-state response (ASSR) is also clear around 40.8 Hz. The average response from 42 NH listeners shows monotonically increasing phase-locking strength of the ITD-evoked response across the ITD magnitudes (Figure 1 C), confirming that the response is indeed sensitive to TFS processing and the size of the ITD jump. Perhaps more important for the search of candidate TFS processing assays, large individual differences are apparent in the phase-locking strength across subjects (Figure 4). Most subjects did not show a salient response for the 20 *μs* condition, and only about half showed robust responses for the 60 *μs* condition. Focusing therefore on the 180 *μs* and 540 *μs* conditions, the 180 *μs* condition is still part of the increasing slope of the response-vs-ITD-jump-size trend, but the response amplitude may have saturated for the 540 *μs*. Accordingly, we used each individual’s response for the 180 *μs* condition for comparison to behavior. Note that the ITD being referred to here is the size of the jump; for instance, for the 20 *μs* condition, the stimulus started with an ITD of 10 *μs* with one ear leading and jumped to the other side about halfway through the stimulus to end with a 10-*μs* ITD with the other ear leading.

**Figure 4.**
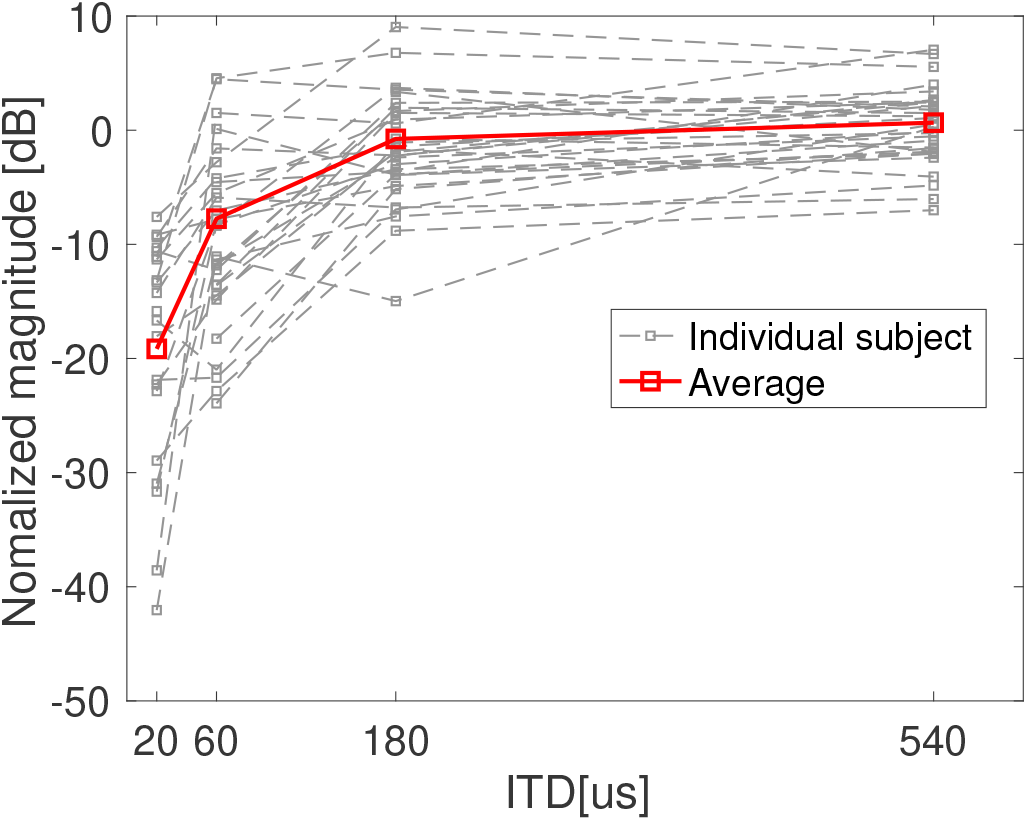
Individual EEG inter-trial coherence (ITC) (averaged under 20 Hz) values as a function of the size of the ITD jump. The ITC increases with the ITD for all subjects. Robust responses above noise floor are detected for most subjects for the “ITD = 180 *μs*” condition. Interestingly, individual differences present at 180*μs* persist even at 540 *μs* despite the ITD jump being obviously perceptible and the response amplitude appearing to saturate.

Unfortunately, one striking aspect of the result in Figure 4 is that even at 540 *μs*, the individual differences that were present in the lower ITD conditions persist. The ITD jump is obviously perceptible at 540 *μs*, and the EEG response appears to be near saturation level for most individuals; this suggests that a significant portion of the individual differences in the magnitude of the cortical response arises from factors extraneous to TFS-based processing. Extraneous factor that may contribute include anatomical factors such as head size, and the geometry/orientation of the neural sources relative to the scalp sensors (Bharadwaj et al., 2019). Thus, although the cortical response to ITD jumps is indeed elicited and parametrically modulated by TFS-based processing, raw response amplitude metrics may be unsuitable for use as an individualized assay of TFS coding.

### 3.3 Candidate subcortical correlates of TFS processing

Figure 5 shows an example FFR recording from a single individual in response to the stimulus sequence with a probe tone, a forward masker, and a second probe tone. The top row (green traces) shows the differential response across two stimulus polarities. This response to the probe tone (labeled “d1” in Figure 5) tracks the 500 Hz TFS in the stimulus, but contains both pre-neural (e.g., cochlear microphonic) and neural components. Because forward masking is thought to arise from synaptic processing (Verschooten and Joris, 2014), the forward masker would be expected to only suppress the neural (i.e., postsynaptic) component of the response to the second probe tone leaving the pre-neural component intact (labeled “d2” in Figure 5). Thus, subtracting d2 from d1 should leave a purely neural response phase-locked to the TFS.

**Figure 5.**
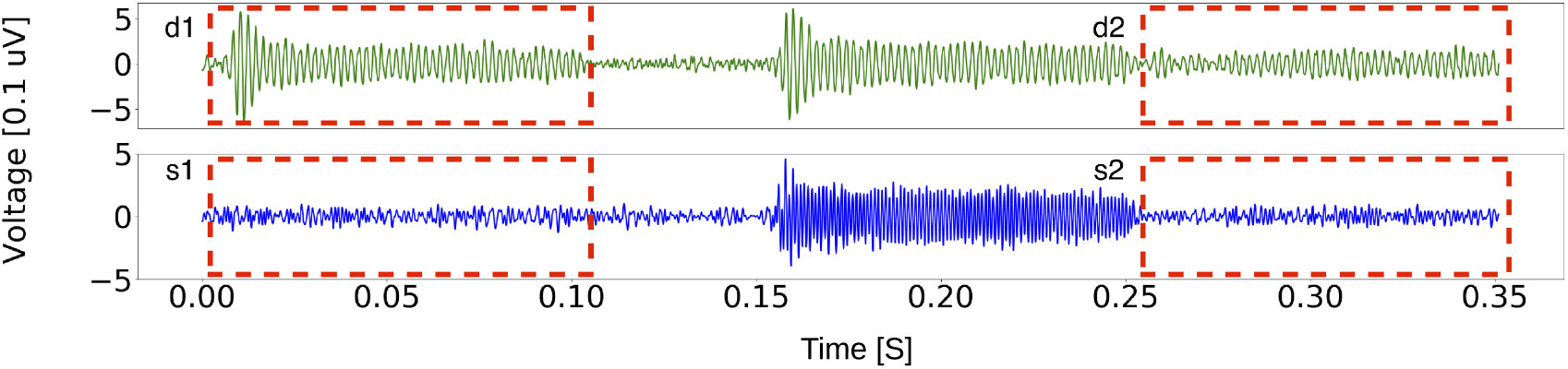
Frequency following response (FFR) to the probe-forward-masker-probe stimulus sequence for an individual subject. The top row (green trace) represents the differential response across two stimulus polarities, whereas the bottom row (blue trace) represents the summed response across two stimulus polarities. The first boxed segments in both rows (red, dashed box, labeled d1 or s1) reflect the raw response to the probe tone, which is likely a mixture of neural and pre-neural responses (e.g., cochlear microphonic; CM), whereas the second boxed segments in both rows (red, dashed box, labeled d2 or s2) is the adapted response after forward masking. For d2 and s2, the pre-neural (e.g., CM) component is expected to be intact whereas the neural response is attenuation by forward masking (only 1-ms gap). The forward masker only partially suppresses the responses, suggesting a strong pre-neural contribution to d1 and s1. The weaker residuals obtained by subtraction, i.e., (d1 - d2) and (s1 - s2) is likely a purely neural.

The bottom row in Figure 5 (blue traces) shows the summed response across the two polarities. Because of inner hair-cell rectification, this response contains a 1000 Hz component arising from the stimulus TFS (also see “Materials and Methods” section). This 1000 Hz component in response to the probe (labeled “s1” in Figure 5) has previously been interpreted as a neural response (Parthasarathy et al., 2020). If that were indeed the case, the forward-masker would considerably suppress the 1000 Hz component in response to the second probe (labeled “s2” in Figure 5).

Figure 6 shows the average d1 (panel A), d1-d2 (panel B), s1 (panel C), and s1-s2 (panel D) response obtained across subjects, quantified in the frequency domain. It is evident from the reduced size of the (d1-d2) response compared to the d1 response, and the reduced size of the (s1 - s2) response compared to the s1 response that, forward masker only has a partially suppressing effect. This provides evidence that both candidate TFS measures—the 500 Hz component from the difference across stimulus polarities, and the 1000 Hz component from the sum across stimulus polarities—have significant pre-neural contributions. This is in contrast to the previous interpretation that the component at double the tone frequency is purely neural (Parthasarathy et al., 2020).

**Figure 6.**
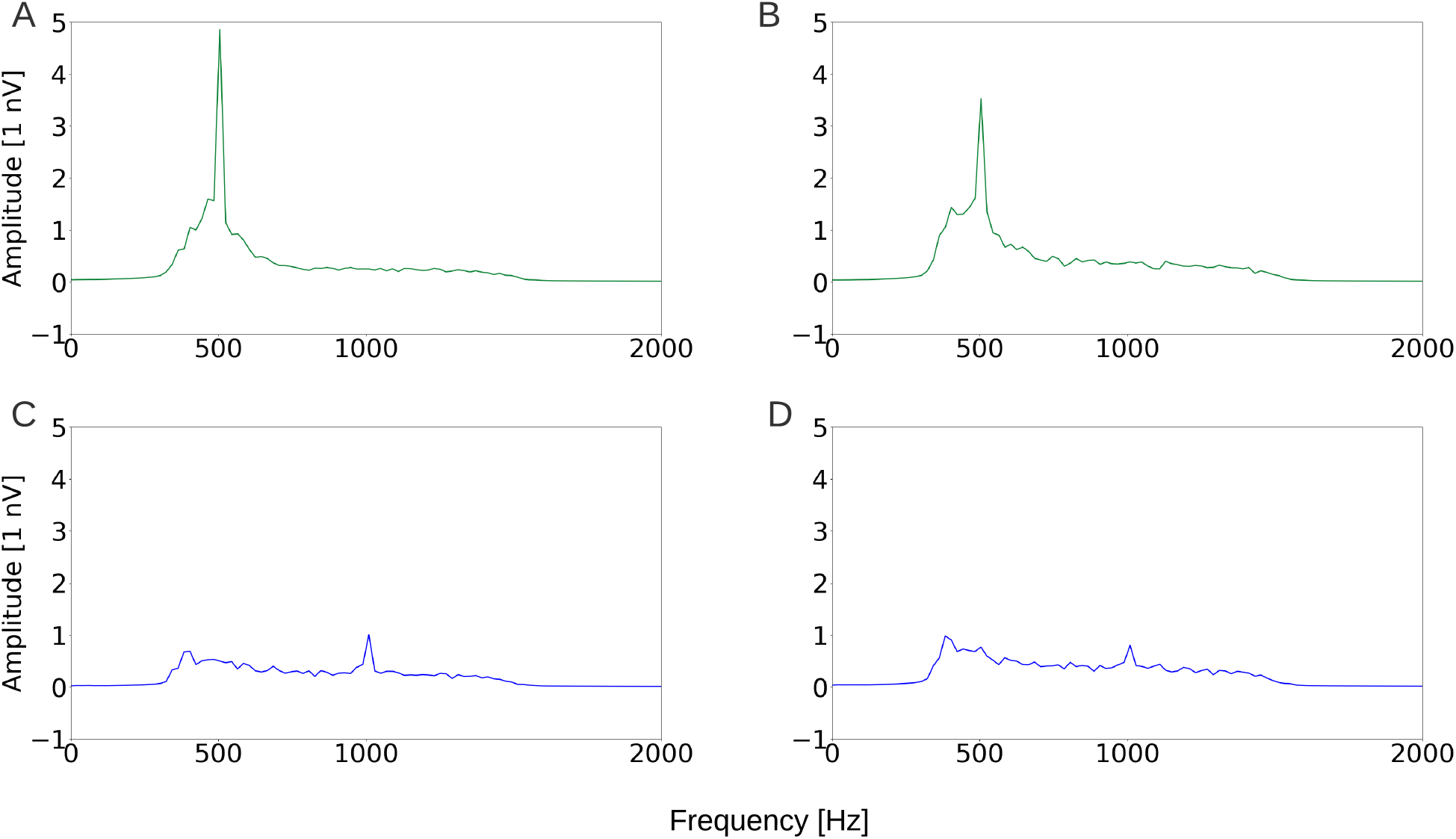
Frequency-domain representations of the d1 (A), d1-d2 (B), s1 (C), and s1-s2 (D) segments from Figure 5, but averaged across subjects. Forward masking partially attenuates both the 500 Hz component of d1 response, and the 1000 Hz component of the s1 response, suggesting that both responses reflect a mix of pre-neural and neural sources.

These results indicate that a forward-masking paradigm will need to be employed to extract the purely neural “residual” response. Unfortunately, unlike transtympanic recordings that are difficult to perform (Verschooten et al., 2015), this residual is small and not readily measurable from all individual subjects. Thus, while subcortical envelope-following responses (EFRs) provide a robustly measurable correlate of envelope processing (Bharadwaj et al., 2015), FFRs tracking the TFS are not promising, and not readily measured across all individuals despite our cohort being comprised of NH listeners.

### 3.4 “Adjusted” behavioral and cortical measures are strongly correlated, likely reflecting TFS coding

Based on individual differences in the cortical amplitude measure persisting for the large-ITD-jump (540*μs*) condition, we concluded that the amplitude measure of cortical phase-locking was dominated by extraneous variance, likely from anatomic factors. Thus, we focused our attention on the latency of the ITD-jump response, because the latency is expected to be unaffected by the scaling effects of individual anatomy. In particular, we extracted the latency of the cortical response to the 180 *μs* jump condition to avoid floor and ceiling effects. The use of the latency metric was also motivated by the previous successful use of this EEG latency measure to predict individual behavioral measures of spatial release from masking (Papesh et al., 2017). In addition to this latency metric, the slope of the cortical response amplitude with increasing ITD-jump (i.e., the increase from the 60 *μs* condition to the 180 *μs* condition, divided by the 540 *μs* condition) was extracted as a normalized measure of TFS processing that would mitigate the overall scaling influence of anatomical factors. This normalization was also motivated by the previous successful use of a similarly normalized electrophysiological measure in the context of modulation processing (Bharadwaj et al., 2015).

Both of these “adjusted” cortical measures exhibited significant correlations with behaviorally measured ITD thresholds. Specifically, individual differences in latency of the cortical ITD-jump response (for 180*μs*) correlated with individual differences in the ITD detection thresholds (R = 0.35, P = 0.048, n = 32). The correlation improved when the behavioral scores were also adjusted to factor out the “nonsensory score” (R = 0.45, P = 0.01, n = 32). The slope metric from the cortical EEG response also correlated with ITD thresholds both with and without adjustments to the behavioral scores (R = 0.43, P = 0.021, n = 32, original ITD scores; R = 0.42, P = 0.028, with “non-sensory score” factored out).

With the subcortical measures, because results indicated a significant pre-neural contribution for both candidate TFS measures, and the residual neural component extracted from the forward-masking paradigm was not robustly measurable for many participants, we did not explore FFR-behavior associations in detail. A simple correlational analysis between the residual (d1 - d2) 500 Hz response and ITD thresholds suggested that the correlations were not statistically distinguishable from zero (not shown).

A multiple linear regression model was used to predict ITD detection thresholds using both the “non-sensory score”, the EEG latency (cortical ITD-jump response), as well as the EEG normalized slope metric. The model could predict the behavioral ITD threshold well (Figure 7) with the predictors together accounting for more than half of the variance observed in the behavioral thresholds (Table 1). We interpreted this result as suggesting that both “adjusted” behavioral scores, and electrophysiological latency or slope metrics in response to TFS-based binaural processing are promising candidate assays of TFS processing that may be suitable for use at the individual level.

**Figure 7.**
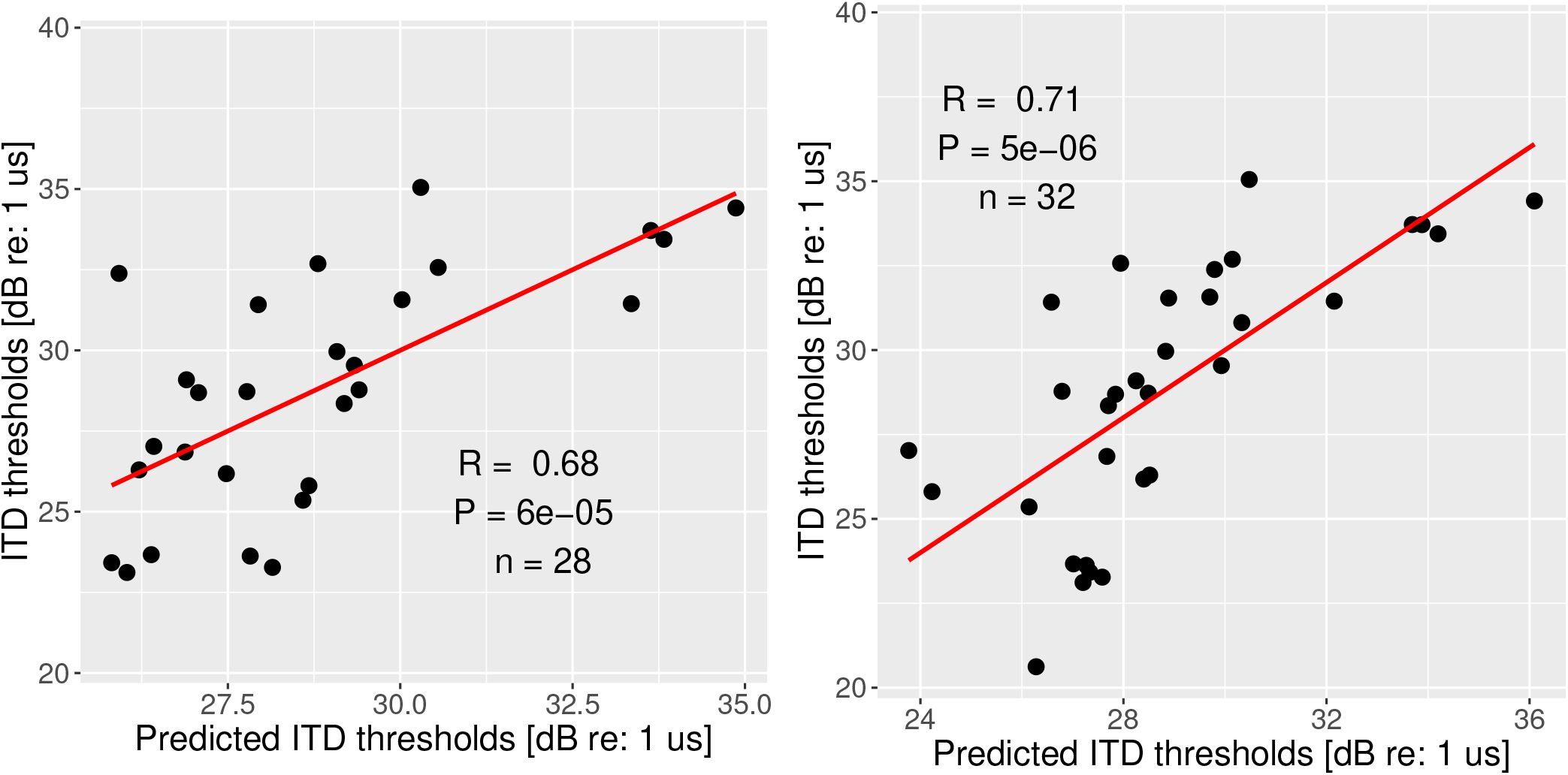
Model prediction of the ITD detection thresholds, based on the combination of lapse rate and slope (60-*μs* to 180-*μs* condition) [left], or the combination of lapse rate and EEG latency [right]. Please refer to Table 1 for the variance explained by each factor.

**Figure 8.**
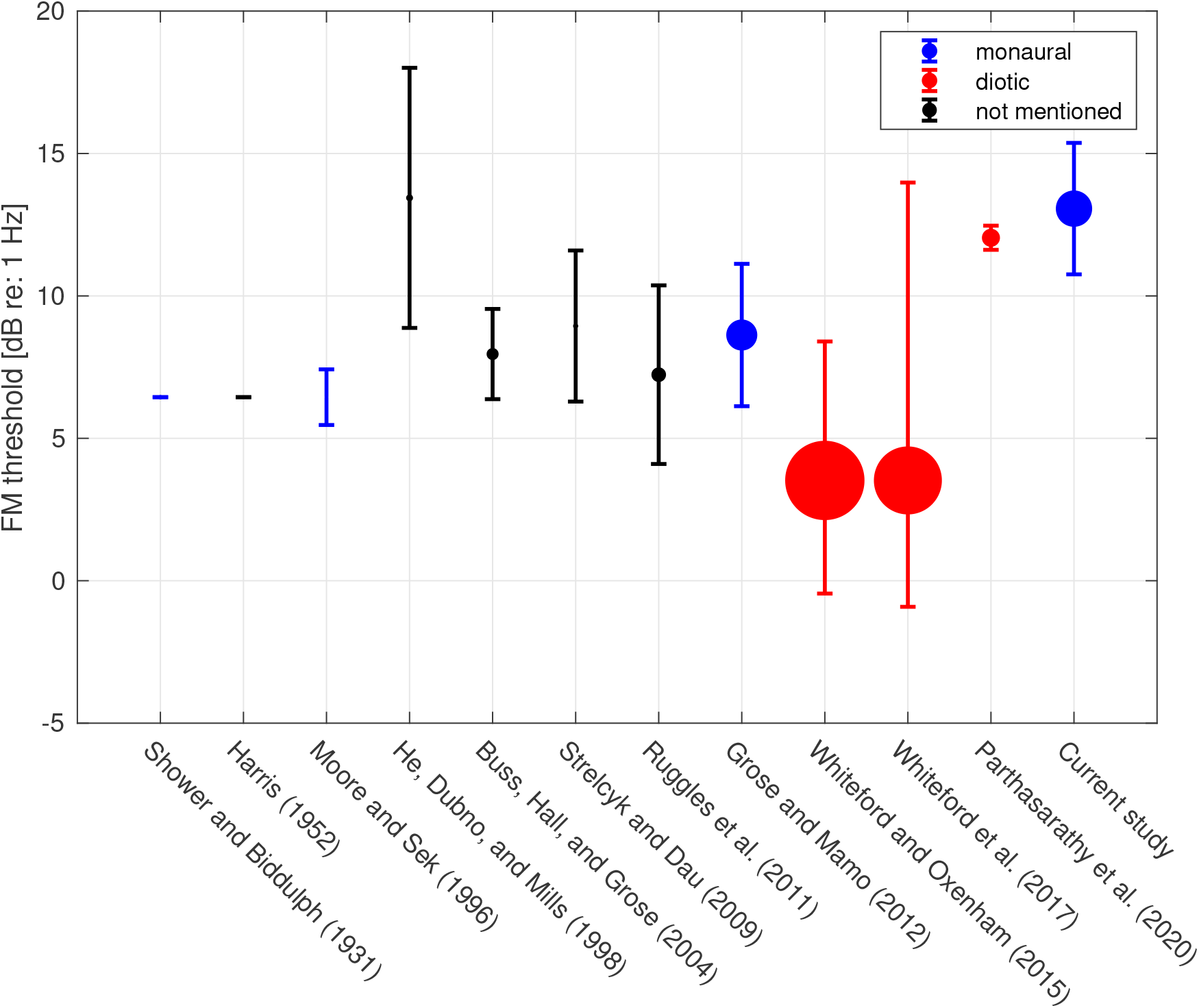
A sample of published reports of FM detection thresholds for comparison (Shower and Biddulph, 1931; Harris, 1952; Moore and Sek, 1996; He et al., 1998; Buss et al., 2004; Strelcyk and Dau, 2009; Ruggles et al., 2011; Grose and Mamo, 2012; Whiteford and Oxenham, 2015; Whiteford et al., 2017; Parthasarathy et al., 2020). Error bar is 1 standard deviation (STD). The size of the dot represents the number of subjects (Whiteford and Oxenham (2015) has the most subjects; N=100). Stimulus parameters such as stimulus level, carrier frequency, and modulation frequency in the cited studies are similar to those used in the current study, with slight differences (e.g., Ruggles et al. (2011); Strelcyk and Dau (2009) used carrier at 750 Hz). Some threshold values are approximate from figures (e.g., mean and std had to be estimated based on median and range in the box whisker plots in Whiteford and Oxenham (2015) and Whiteford et al. (2017)). The mean and std from the young and middle-aged group from Grose and Mamo (2012) were combined to generate a single data point. Some authors expressed the threshold in terms of Δ*F/F*_*c*_, where Δ*F* is frequency deviation, and *F*_*c*_ is the carrier frequency. Moore and Sek (1996) used Δ*F* that was in two directions, i.e., peak-peak. Subjects from some studies were highly experienced in psychoacoustic tasks hence the thresholds were very low/good. Whiteford and Oxenham (2015); Whiteford et al. (2017) obtained thresholds that fall in the lower end of the results of the current study from a very large number of subjects. This may be because their subjects were younger normal-hearing listeners and the stimuli were presented diotically and dichotically instead of monaurally.

**Figure 9.**
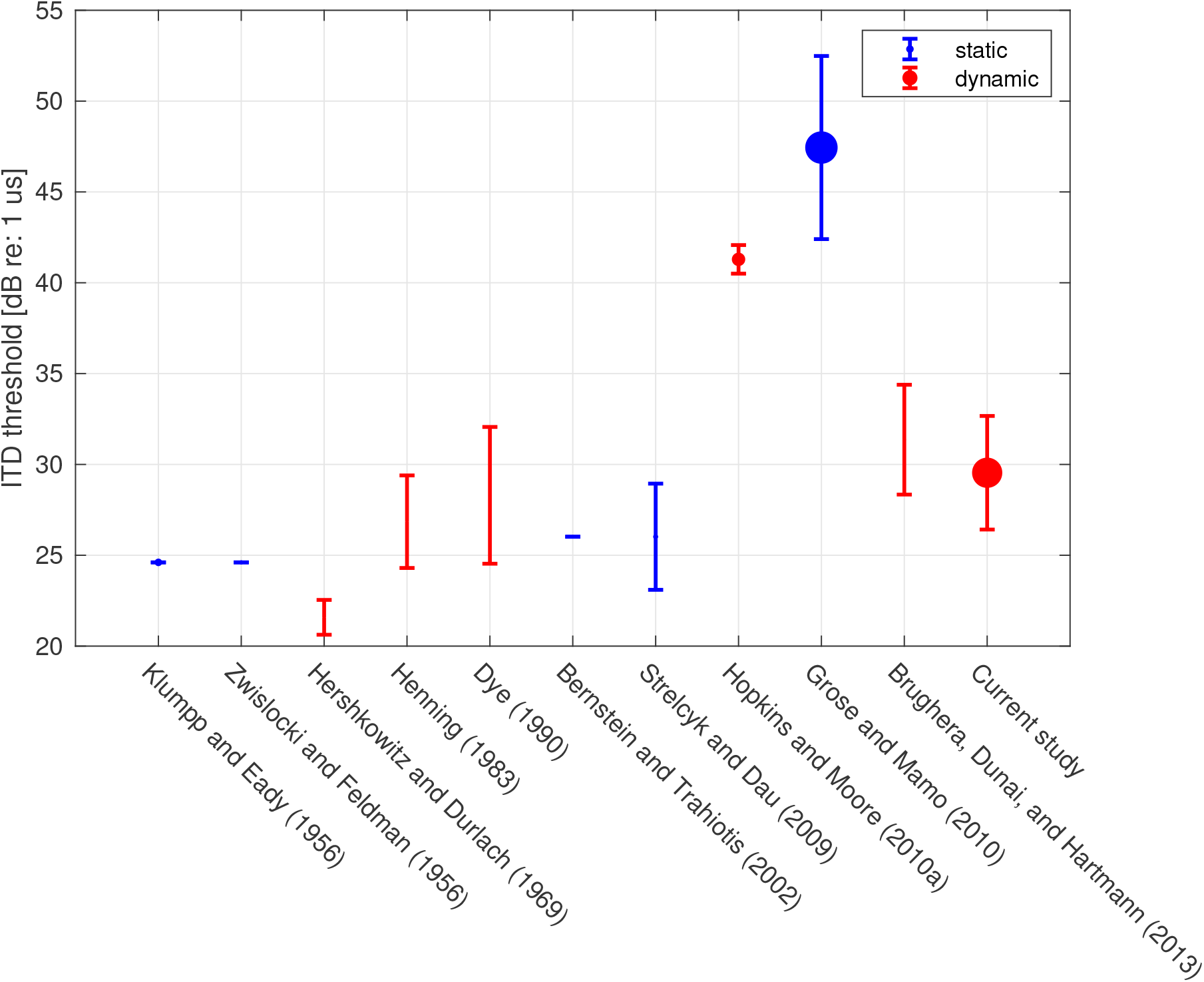
A sample of published reports of ITD detection thresholds for comparison (Klumpp and Eady, 1956; Zwicker, 1956; Hershkowitz and Durlach, 1969; Henning, 1983; Dye, 1990; Bernstein and Trahiotis, 2002; Strelcyk and Dau, 2009; Hopkins and Moore, 2010; Grose and Mamo, 2010; Brughera et al., 2013). Error bar is 1 standard deviation (std). The size of the dot represents the number of subjects (the current study has the most subjects; N=36). Stimulus parameters such as level and carrier frequency in the cited studies are similar to those used in the current study, with slight differences (e.g., Strelcyk and Dau (2009) used carrier at 750 Hz). Note that some threshold values were extracted approximately from figures rather than direct numerical reports. Some of the studies used stimuli with the leading ear switching from one size to the other (labeled “dynamic”, marked in red color), whereas others presented an ITD only in the target intervals, with the reference being the midline (labeled “static”, marked in red color). Note that the values from Hershkowitz and Durlach (1969); Brughera et al. (2013) were halved since the authors used *ITD/*2 in each interval. The mean and std from young and middle-aged cohort from Grose and Mamo (2010) were combined to generate a single data point. Subjects from some studies were highly experienced in psychoacoustic tasks.

**Table 1.** Model prediction of the behavioral ITD detection thresholds, with factors including the “non-sensory score”, EEG latency, and EEG slope. The variations accounted for by the “non-sensory score” are more than three times as by either one of the two EEG metrics. Together, more than half the variance is accounted for.

| Predictor | Variance Explained |
| --- | --- |
| Non-sensory score | 37.48% |
| EEG latency | 10.03% |
| EEG slope | 9.28% |
| Explained | 56.79% |
| Unexplained | 43.21% |

## 4 Discussion

In the present study, we sought to identify viable assays that can index the fidelity of TFS processing at the individual subject level. To obtain insight into whether individual differences in various candidate measures reflected TFS-based processing or extraneous factors, we compared individual differences in behavioral scores across FM and ITD detection tasks to differences in cortical and subcortical EEG-based measures. Results revealed the strong influence of extraneous factors on both behavioral scores and amplitude-based EEG metrics.

With behavioral measures, non-sensory factors, quantified using the lapse rate in catch trials, could account for a third of the variance across individuals. Although previous work has explored a range of behavioral TFS measures (Moore and Sek, 2009; Hopkins and Moore, 2010; Sek and Moore, 2012), the results from the present study underscore the importance of adjusting raw behavioral scores to reduce the impact of non-sensory factors. Indeed, although raw FM and ITD measures correlated significantly with each other, similar to the correlation between monaural AM detection and binaural envelope-ITD thresholds (Bharadwaj et al., 2015), this was driven in part by non-sensory factors. Because phase-locking to the TFS is essential for low-frequency ITD processing (Yin and Chan, 1990), it is plausible that ITD detection thresholds can provide an index of TFS sensitivity. On the other hand, whether FM detection relies on TFS coding has been controversial due to the possible role of recovered ENV cues that result from cochlear filtering of FM stimuli; indeed, FM stimuli lead to perceptible out-of-phase ENV fluctuations at cochlear places tuned to frequencies just above and below the FM carrier (Whiteford et al., 2017; Whiteford and Oxenham, 2015). Whiteford et al. (2020) extensively tested the role of place coding in FM detection and found that place coding by itself can account for the observed variations in FM sensitivity across all carrier frequencies and modulation rates. This finding is in contrast to the widely accepted view of the utilization of time coding in the detection of slow-rate FM (Strelcyk and Dau, 2009; Moore and Sek, 1996; Parthasarathy et al., 2020). Together with our finding that non-sensory factors influence raw behavioral scores, this uncertainty about the link between TFS coding and FM detection calls into question the previous use of FM detection scores as a correlate of TFS processing. In contrast, unambiguous theoretical links can be made between ITD detection and TFS coding, suggesting that once ITD thresholds are adjusted to reduce the influence of non-sensory scores, they may serve as a useful metric of TFS processing. This was corroborated by our finding that passive EEG measures, when combined with non-sensory scores, can account for more than half the variance in ITD thresholds. Here, we used lapse rates in the catch trials to obtain a correlate of non-sensory factors. Alternately, a surrogate behavioral task that does not rely on TFS coding (e.g., interaural level difference sensitivity) may also be used to adjust ITD thresholds with similar benefits.

Another key finding from the present study is that although passive EEG measurements can potentially reflect TFS-based processing objectively, they too are susceptible to the influence of extraneous factors. Indeed, consistent with the interpretation that individual anatomical factors can have a scaling influence on response amplitudes, we found that cortical responses phase-locked to ITD changes showed large individual differences even for a large ITD jump (540 *μs*) where the response amplitude was near saturation for most individuals. Therefore, we argued that the evoked-response latency and/or percent growth/slope metrics may be better assays of TFS processing. Accordingly, latency and slope metrics showed significant correlations with behavioral ITD detection thresholds. For candidate subcortical FFR-based measures of TFS processing, our results showed that pre-neural physiological currents (cochlear microphonic, inner hair-cell currents) contribute significantly to the measure, thus complicating their applicability. Indeed, brainstem response measures from individuals with compromised inner hair-cell synaptic transmission show that pre-neural transduction currents can contribute to the measured response (Santarelli et al., 2009). Moreover, when employing a forward-masking-based design to isolate the neural component of the FFR, the resulting signal is relatively weak even in our NH cohort. This result from non-invasive ear-canal recordings is in contrast to neurophonic measurements from the auditory nerve (Snyder and Schreiner, 1985) or round window (Henry, 1995) from animals, or FFR measurements from humans using transtympanic electrodes where the forward-masking design has been used successfully (Verschooten et al., 2018). Although FFRs have previously been used as a putative correlate of TFS-based processing (Parthasarathy et al., 2020), our results suggest that further experiments are needed to clarify the interpretation of those results and to enhance the quality of the measured signal across.

Our finding that the FFR may be a poor correlate of neural TFS processing is in contrast to previous results using subcortical envelope-following responses (EFRs) are correlates of ENV processing. For example, Bharadwaj et al. (2015) showed that the AM detection thresholds and ENV-based ITD thresholds correlated strongly with normalized EFR-based metrics. This is likely both because EFR measurements more readily exclude pre-neural contributions (which primarily track the TFS), and because Bharadwaj et al. (2015) obtained asymptotic behavioral scores from a large number of trials (1200-1500 trials) from trained subjects. Indeed, with naïve subjects in this study, an AM detection task similar to the one used in Bharadwaj et al. (2015) also showed a strong influence of non-sensory factors.

In summary, the present study examined various candidate assays for quantifying TFS processing at the individual subject level. These included behavioral FM and ITD detection thresholds, EEG-based cortical and subcortical physiological measures. Among these, our experiments suggest that the latency of cortical responses to ITD jumps, normalized cortical response amplitude (i.e., percent growth/slope), and “adjusted” ITD thresholds may all be useful. Indeed, when a multiple linear regression model was constructed to predict behavioral ITD thresholds, the combination of the non-sensory score (lapse rate in catch trials), EEG latency, and slope measures could account for more than 50% of the variance across individuals. Our results are consistent with the findings by Papesh et al. (2017), who also found a correlation between ITD-evoked EEG latency and spatial-hearing outcomes such as spatial release from masking. Given that multiple candidate measures were explored to identify the most promising assays, future experiments should be conducted to independently confirm the efficacy of the assays endorsed by our results. The most promising assays rely on binaural TFS-based processing.

Reliable measures of TFS processing are critical for future investigations into the role of the TFS in everyday hearing using intact speech-in-noise stimuli without vocoding manipulations. While sub-band vocoding can allow for independent manipulation of acoustic TFS and envelope cues, subsequent cochlear processing can confound these factors once again (Gilbert and Lorenzi, 2006; Swaminathan and Heinz, 2012). Furthermore, when both rate-place/ENV cues and TFS cues are redundant information, vocoding experiments cannot provide insight into how are perceptually weighted. The candidate TFS measures identified in the present study can help address these gaps.

## Notes

### Competing Interest Statement

The authors have declared no competing interest.

